# Effects of plant evolution on nutrient cycling couple aboveground and belowground processes

**DOI:** 10.1101/071126

**Authors:** Nicolas Loeuille, Tiphaine Le Mao, Sébastien Barot

## Abstract

Plant strategies for nutrient acquisition and recycling are key components of ecosystem functioning. How the evolution of such strategies modifies ecosystem functioning and services is still not well understood. In the present work, we aim at understanding how the evolution of different phenotypic traits link aboveground and belowground processes, thereby affecting the functioning of the ecosystem at different scales and in different realms. Using a simple model, we follow the dynamics of a limiting nutrient inside an ecosystem. Considering trade-offs between aboveground and belowground functional traits, we study the effects of the evolution of such strategies on ecosystem properties (amount of mineral nutrient, total plant biomass, dead organic matter and primary productivity) and whether such properties are maximized. Our results show that when evolution leads to a stable outcome, it minimizes the quantity of nutrient available (following Tilman's R* rule). We also show that considering the evolution of aboveground and belowground functional traits simultaneously, total plant biomass and primary productivity are not necessarily maximized through evolution. The coupling of aboveground and belowground processes through evolution may largely diminish predicted standing biomass and productivity (extinction may even occur), and impact the evolutionary resilience (ie, the return time to previous phenotypic states) of the ecosystem in face of external disturbances. We show that changes in plant biomass and their effects on evolutionary change can be understood by accounting for the links between nutrient uptake and mineralization, and for indirect effects of nutrient uptake on the amount of detritus in the system.

## Introduction

Nutrient cycling is a key component of ecosystem functioning. It is strongly influential for primary production and exerts a bottom-up control on the composition of food webs (i.e., primary producers, herbivores, predators) (Vitousek et al. 1997). Numerous plant traits (e.g. nutrient uptake rate, biomass turnover, litter quality and influence on mineralization through rhizosphere priming effect) influence the intensity of nutrient cycling rates (Chapin III et al. 2002),. Such traits directly affect aboveground and belowground processes. For instance, nutrient uptake rate and biomass turnover constrain aboveground biomass, while plant control on mineralization can change belowground characteristics such as nitrogen or carbon contents.

From a functional point of view (Chapin III et al. 2002) and, more recently from an evolutionary point of view (Loeuille et al. 2002; Loeuille and Loreau 2004; Loeuille and Leibold 2008; Boudsocq et al. 2011), the links between nutrient uptake rate and plant individual biomass turnover have been largely investigated. For example, to take up more mineral nutrient plants may produce more thin and short-lived roots or sustain a large mycorrhizal network, providing organic matter in exchange for mineral nutrients. Such strategies incur allocation costs, diverting energy from plant individual growth or reproduction (e.g., Cheng & Gershenson 2007). Such allocation costs explicitly link aboveground (plant individual growth) and belowground (mineralization activation) processes. The novelty of the present work lies in the investigation of how such a link affects the evolution of plant strategies and ecosystem functioning.

Considering such a coupling, evolution of plant traits simultaneously affects food webs that are often separated, i.e. belowground and aboveground food webs. Reciprocal effects between aboveground and belowground topic currently raises increasing interest (Zou et al. 2016) and the plant compartment is central in understanding this interaction. Evolutionary dynamics may lead to contrasted outcomes regarding the quantities 59 of nutrient stocked aboveground (proportional to total plant biomass) vs belowground (detritus) with important consequences for the global dynamics of ecosystems. While total plant biomass determines the amount of energy available for higher trophic levels aboveground, the amount of detritus influences the total energy available to belowground detritivore food webs. In turn, available energy largely impacts the length of food chains (Oksanen et al. 1981; Loeuille and Loreau 2005) and food web stability (Rosenzweig 1971). Evolutionary dynamics associated with these traits thus have far reaching implications.

Our goal is to go beyond the traditional focus of evolutionary functional models modelling plant growth and mortality traits, by linking such traits to belowground processes such as mineralization. We model the evolution of nutrient uptake rate, and its consequences for nutrient turnover and mineralization due to allocation trade-offs. We then assess the evolutionary consequences for ecosystem properties. The evolutionary outcome critically depends on the shape of the trade-off functions, but we only find three qualitatively different ecological outcomes: extinction of the plant population, continuous accumulation of nutrient during evolution, or evolution toward stable ecosystem properties. While the ecological model is based on a previous article (Boudsocq et al. 2011), our approach is novel in at least two ways. First, it focuses on different traits, with an explicit focus on mineralization, thereby linking evolution to nutrient acquisition and retention explicitly. This allows a coupling between aboveground and belowground processes, providing a more integrative view of eco evolutionary dynamics of plant strategies. Second, by considering that evolution involves existing links between four different traits (basic growth rate, competitive ability, nutrient turnover and mineralization), while Boudsocq *et al*. (2011) (and most evolutionary models in ecology) couple only two traits in trade-off functions. The multi-dimensionality of evolutionary dynamics is a rising and important question in evolutionary ecology (Gilman et al. 2012) and we hope that our work may help to understand its implications 84 for the evolution of plant strategies.

We focus on a restricted number of issues: How is the phenotypic composition of the plant community modified through evolution? What are the ecosystem properties associated to these evolutionary outcomes (amount of mineral nutrient, total plant biomass, dead organic matter and primary productivity)? Are these properties maximized as a result of evolution? We show that coupling aboveground and belowground processes strongly modifies predicted dynamics, even in the case of the non-spatial model we employ here. The coupling can enhance or reduce predicted standing biomass and productivity, affecting the evolutionary resilience (i.e., the time it takes for evolutionary dynamics to go back to the selected strategy) in the face of environmental perturbations (such as climate change, increase of fertilizers, fires, erosion).

### Methods

We model the dynamics of a limiting nutrient inside an ecosystem composed of three compartments: inorganic nutrient (*N*), plants (*P*) and dead organic matter (*D*) (Figure 1). *N*, *P* and *D* correspond to the quantity of limiting nutrient in each compartment (most usually, nitrogen). While compartments are quantified in terms of limiting nutrient, we do not account for plastic or evolutionary variations in stoichiometric ratios or in organism size, thus implicitly assuming them constant, so we refer to *P* as plant biomass hereafter. Time variation of nutrient stocks can be written:

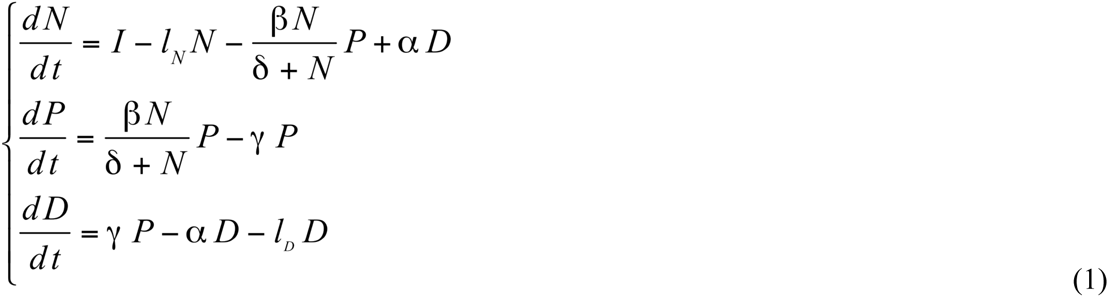

**Figure 1.**
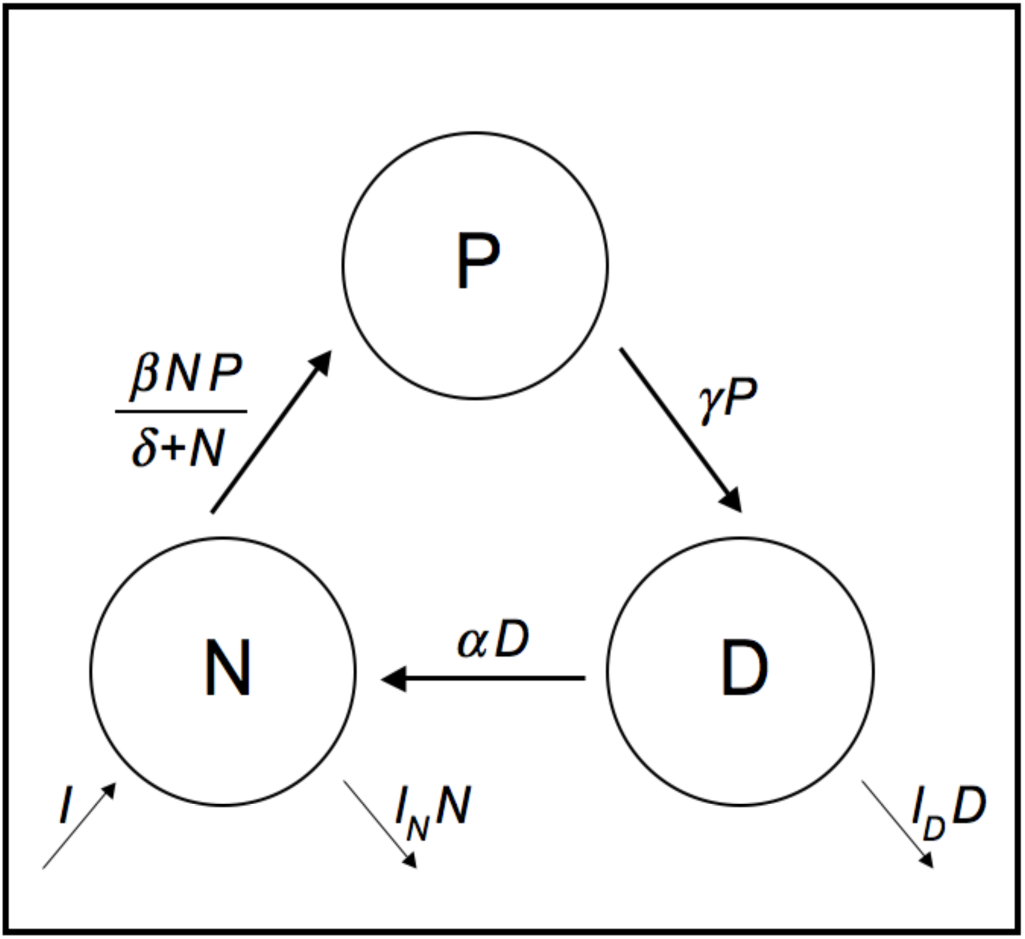
Modeled nutrient compartments and fluxes. Circles represent ecosystem nutrient compartments: inorganic nutrient (*N*), plants (*P*) and dead organic matter (*D*). Arrows and correspond to nutrient fluxes.

Parameters *β* and *δ* define the plant growth rate using a classical Monod function. Primary productivity *ϕ* is defined by the uptake term *βNP/(δ+N)*. Parameter *γ* defines the turnover rate of plant biomass. Through evolution and trade-offs (see below), traits *β*, *γ* and *δ* influence the mineralization rate *α*. The model thus couples aboveground (eg, plant growth/production) and belowground processes (nutrient uptake, mineralization) explicitly. Parameters describing global inputs and outputs of nutrient are *I* and *I_N_*, *I_D_* respectively. The model is simple as it focuses on one plant compartment with one limiting factor (a nutrient). Including other density dependent effects (due to space or light competition) or community aspects (multiple species) would of course make it more realistic. We do not account for such additional components to keep the evolutionary dynamics tractable and focused on existing links between aboveground and belowground processes. For more details on the parameters of the model and parameter values, see Table S1.

### Relation between internal cycling rates - plant strategy trade-offs

The model assumes that different aspects of plant life history–competitive ability, biomass turnover, mineralization-are directly linked to intrinsic growth and reproduction due to allocation constraints. Intrinsic growth and reproduction, being the rate of increase in plant biomass when nutrient is not limiting, corresponds to *β*. Competitive ability, as measured by the rate of growth when nutrient is rare, is directly (and negatively) linked to *δ*. Biomass turnover is proportional to *γ*, and we consider this turnover to be either intrinsic (e.g., root or leaf loss) or due to enemies (herbivores, pathogens, etc.). Mineralization is constrained by parameter *α*;. It embodies both intrinsic properties such as litter degradability and the activation of decomposers (e.g., microbes) by the plant, through the release of activating compounds.

To account for allocation costs, we propose to write parameters describing nutrient uptake and recycling as:(*α*,*δ* or *γ*) = (*k_1_*\* *β*+*k_2_*)^*g*^ (see table 1 for the relationships and their biological justifications). We use such functions because of their flexibility. They may be linear, concave or convex depending on the value of exponent *g*. Such a flexibility is desirable, because the shape of trade-off functions is usually not known empirically, but largely matters for the outcome of evolutionary dynamics (de Mazancourt and Dieckmann 2004; Loeuille and Loreau 2004).

**Table 1:** Biological justification of trade-off functions used in the present work.

| Traits considered | Relationship | Biological justification | References | Function used |
| --- | --- | --- | --- | --- |
| $\delta$ vs $\beta$ | positive | High growth rate when rare incurs a cost in terms of competitive ability | r-K Theory (McArthur and Wilson 1967; Pianka 1970) | $\delta=(a\beta+\delta_0)^{g_1}$<br>(2) |
| $\gamma$ vs $\beta$ | positive | -Fast growing species have high nutrient uptake and large turnover rate<br>-Investment in defenses reduce turnover, but incurs costs in growth or reproduction | Eissenstat *et al.* (2000)<br>Herms & Mattson (1992) | $\gamma=(m\beta+\gamma_0)^{g_2}$<br>(3) |
| $\alpha$ vs $\beta$ | negative | -Activation of decomposers (eg, microbes) increases mineralization $\alpha$ but is costly in terms of growth and reproduction<br>-production of tough tissues reduces mineralization | Cheng & Gershenson (2007)<br>Whitham *et al.* (2003)<br>Grime *et al.* (1996) | $\alpha=(-r\beta+\alpha_0)^{g_3}$<br>(4a) |
| | positive | -For fast growing plant species, increased nutrient content in tissue makes litter more easy to mineralize<br>-Increased carbon input in soil may accelerate litter decomposition (priming effect) | Chapin (1980)<br>Berendse (1994)<br>Fontaine, Mariotti & Abbadie (2003) | $\alpha=(r\beta+\alpha_0)^{g_3}$<br>(4b) |

Because of these trade-off functions, our model links aboveground and belowground processes in a single evolutionary framework. Parameters *β* and *δ* for instance determines the nutrient uptake (belowground), but also the increase in plant biomass (part aboveground, part belowground). Parameter *γ* describes the loss of plant biomass (again, part aboveground, part belowground) to the detritus compartment. Finally, *α* represents the belowground process of mineralization.

### Adaptive dynamics of plant phenotypic traits

We study the evolutionary dynamics of nutrient uptake *β* using the adaptive dynamics methodology (Dieckmann and Law 1996; Geritz et al. 1998). The other traits are deduced from the allocation trade-offs (Table 1). Because these functions are strictly monotonic, choosing another trait as a basis instead of nutrient uptake *β* would produce similar results. Adaptive dynamics model the evolution of phenotypic traits based on clonal reproduction, leaving out the genetic basis, and assuming that evolutionary dynamics are sufficiently slower than ecological dynamics. Although these hypotheses may seem restrictive, they allow a thorough analytical study of selective regimes and of their consequences for ecological systems. Evolution proceeds by the successive replacements of one phenotype by another, a process shown to be similar to expected patterns of trait-based community assembly. While the initial derivation of adaptive dynamics is strongly grounded in evolutionary perspectives,results often extend to other types of adaptation (eg, changes in behaviour, plasticity: Abrams 5). Evolution of nutrient uptake *β* is modeled using the canonical equation of adaptive dynamics:

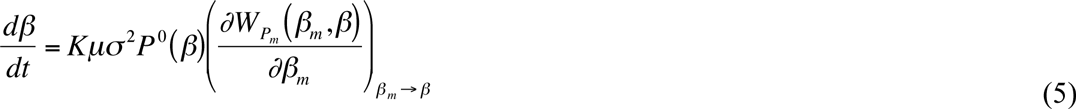

where the fitness of the mutant *β_m_* is deduced from its population dynamics:

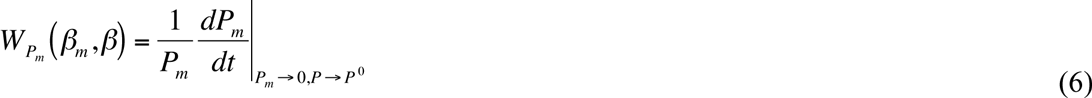

and with *K*: scaling constant, *μ*: per unit biomass mutation rate, *σ*^2^: variance of the amplitude of mutations, *P^0^*: plant biomass at ecological equilibrium.

The selection gradient 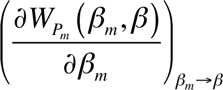determines the direction of evolutionary trajectories. Evolutionary singularities *β*° are obtained for 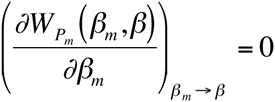. The second derivatives of plant individual fitness with nutrient uptake *β_m_* and *β* give the properties (invasibility and convergence) of evolutionary singularities (Geritz et al. 1998). A singularity is convergent provided:

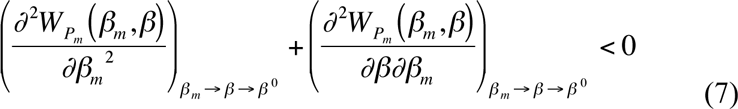

Convergence insures that selection will favor strategies closer to the singularity in its vicinity. The strategy is non-invasible when:

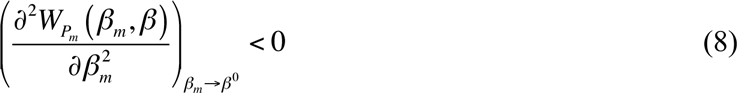

Then, no mutant can invade at the evolutionary singularity. When both equations (7) and (8) are satisfied, *β*° is a continuously stable strategy CSS (Eshel 1983), noted *β_CSS_*. Evolution stops once *β_CSS_* is reached.

Because we have analytical expressions of the ecological equilibrium (*N^0^*,*P^0^*,*D^0^*) it is possible to determine how evolution impacts ecosystem stocks and primary productivity. We compute their derivatives regarding nutrient uptake *β* and combine them with equation (5). Let *X*° denotes one of these variables:

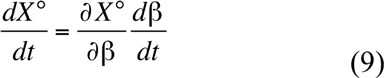

## Results

### Impacts of evolution on system functioning

We here summarize the main results. For detailed information, see appendix 2. Setting equations (1) to zero determines the position of the ecological equilibrium. A unique nontrivial equilibrium exists:

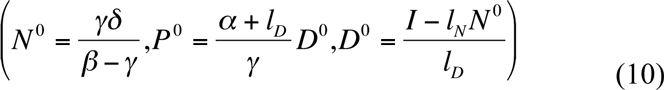

While nutrient uptake *β*, biomass turnover *γ* and competitive ability *δ* influence all three compartments, mineralization α only influences *P^0^*.

Variables *N*, *P* and *D* being positive, it is necessary that:

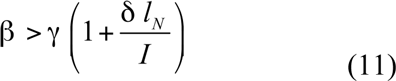

If this condition is satisfied, the equilibrium is also stable.

The fitness of a mutant is:

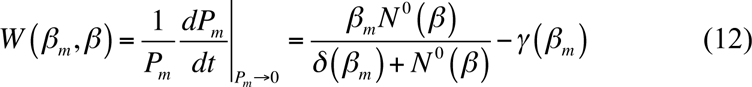

Note that a direct implication of equation (12) is that the mutant can invade (ie, W is positive) if and only if 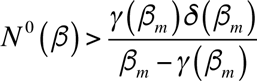 which, given equation 10, can be rewritten 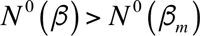. It follows that the mutant can invade only provided it leaves less nutrient at equilibrium than the resident, following Tilman's R* rule (Tilman 1982).

From (12), the selection gradient is:

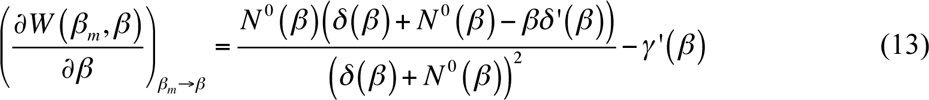

Only two types of evolutionary dynamics can take place. (1) runaway evolution, nutrient uptake *β* being always selected (figure 2a) or counterselected; (2) a *β_CSS_* exists and evolution eventually settles there, provided the CSS allows the existence of the system (ie, it satisfies condition (11) and allows the positivity of equation (4a) (Table 1)). Following Boudsocq *et al*. (2011), we propose to categorize these evolutionary outcomes depending on their consequences for ecosystem functioning (for exact conditions, see appendix 3):

1. “Explosive R* scenarios” (eg, figure 2A). In such scenarios, we have a continuous evolution of traits that leads to ever-increasing plant biomass (hence “explosive”), while mineral nutrient are minimized, in agreement with Tilman's R* rule (hence “R*”). Eventually, crucial hypotheses of the ecological model will be violated, as another constraint (space, light, water, alternative nutrient) will become limiting.
2. “Tragic R* scenarios” (eg, figure 2B). In such scenarios, evolution selects for traits that either continuously erode plant biomass and productivity, or lead the system out of the range of existence (figure 2B), hence the “tragic”. Inorganic nutrient is still minimized (hence “R*”). Note that such scenarios may happen either because runaway evolution continuously erodes plant biomass, such that it may become vulnerable to demographic stochasticity, or because *β_CSS_* falls outside of the range of existence (equations 4a (Table 1) and 11).
3. “Realized R* scenarios” (eg, figure 2c, 2d). In such instances, evolution leads to a stable functional state in which plant biomass and productivity is positive (hence “realized”) while inorganic nutrient is still minimized (hence “R*”). Two situations are then possible: either nutrient uptake *β* always increases through evolution, while nutrient stocks asymptotically tend toward positive values or nutrient uptake *β* eventually settles at a CSS value where ecosystem compartments have positive nutrient mass (figure 2c-d). In both cases, the system reaches a stable and feasible functional state.

**Figure 2:**
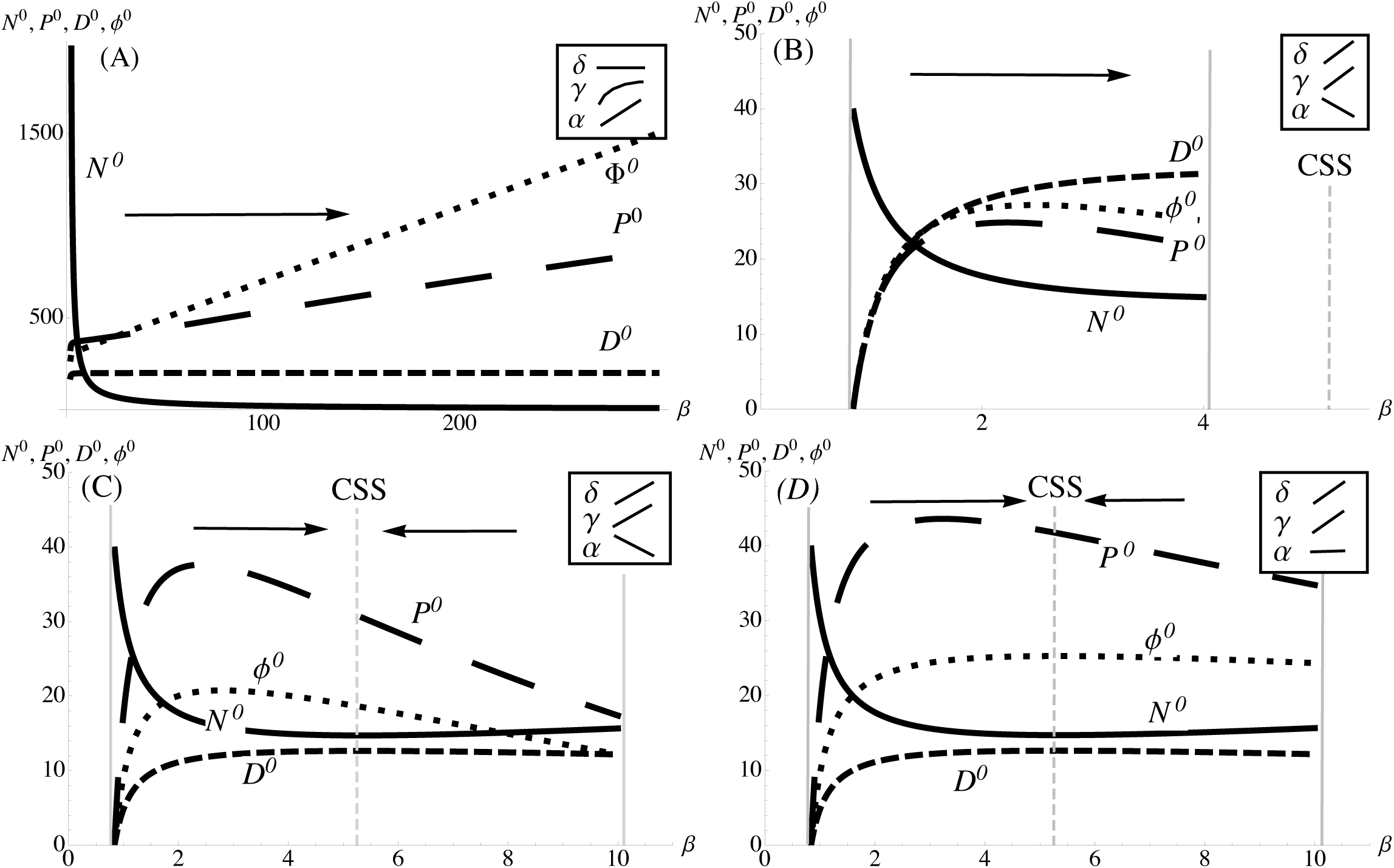
Variation in the size of ecosystem compartments (*N*°,*P*°,*D*°) and primary productivity (*ϕ*°) depending on trait *β*. *N*°: solid line, *D*°: dashed line, *P*°: Long-dashed line, (*ϕ*°: dotted line. Arrows indicate the direction of evolution. Vertical solid grey lines show the boundary of existence for the model. Grey dashed vertical lines indicate the value of the selected strategy. Insets indicate variations of *α*,*β*,*γ* with *β* (trade-off functions, see table 1). a: “Explosive R* strategy”, while increasing *β* is selected, plant biomass and primary productivity increase continuously. Compartments and productivity are rescaled: *N*° *(x1000)*, *D*° *(x50)*, *ϕ(x0.5)*. b: “Tragic R* strategy”, *β* converges to *α*_0_/*r*, at which point rate mineralization is null. Compartments and productivity are rescaled: *N*° *(x10)*, *D*° *(x25)*, *ϕ(x2)*. c-d: “Realized R* strategy” *β* converges to the selected strategy. Compartments and productivity are rescaled: *N*° *(x10)*, *D*\* *(x10)*. c: no maximization of the primary productivity. d: maximization of the primary productivity.

The examples shown on figure 2 can give some insights regarding the mechanisms at hand for falling in one or another category (see also supplementary information for more general results). The shape of trade-off functions is particularly critical in this regard. Consider fitness gradient (13). It clearly underlies the crucial role of variations in biomass turnover *γ* and competitive ability *δ* with nutrient uptake *β* as constraints for the direction of evolution. If the costs in terms of competitive ability (increasing *δ*) or in terms of biomass turnover (increasing *γ*) are not strong (constant or concave functions, figure 2a, see also supplementary information), evolution of ever-increasing nutrient consumption *β* is predicted. Such an increase in nutrient uptake *β* can either lead to explosive R* (on the condition that *P^0^* continuously increases when nutrient uptake *β* increases, ie, mineralization *α* increases faster than biomass turnover *γ* with nutrient uptake *β*), or to a tragic R* (when, conversely, *P^0^* is negatively affected by increases in nutrient uptake *β*). On the contrary, when evolution of *β* is quite costly (ie, competitive ability *δ* or biomass turnover *γ* varies in a linear or convex fashion with *β*), then a selected strategy (CSS) exists (figure 2b-d). The position of such a selected strategy may be outside the range of existence of the system, a “tragic R*” scenario (figure 2b). However, increasing basic mineralization (figure 2b vs 2c,d) enlarge the range of existence and allows a realized R* scenario (figure 2c,d).

### Are the functional properties (nutrient stocks and primary productivity) maximized through evolution?

Because evolution is based on individual fitness (equation 12), links with emergent ecosystem properties can only be indirect. *A priori*, there is no reason to expect that evolution optimizes the system in any way. Evolution however leads to systematic variations in the compartments and fluxes within the ecosystem, depending on the evolutionary scenario.

For “Explosive R* strategies” (Figure 2a), standing plant biomass increases by definition through evolution. Primary productivity also increases. The quantity of inorganic nutrient is minimized while the dead organic matter compartment is maximized. Higher nutrient input (*I*) or lower detritus outputs (*l_D_*), increase the detritus compartment, plant biomass and productivity. This global redistribution of nutrient, from the inorganic compartment to the other compartments can be explained again from trade-off shapes. Because loss terms are bounded (*γ* is of concave shape), and because mineralization *α* increases with the evolution of higher nutrient uptake *β*, plants acquire increasing amounts of nutrient. In the case of “Realized R* strategies” contrasted outcomes are possible. In runaway evolution instances, inorganic nutrient is minimized, plant biomass, primary productivity and dead organic matter are all maximized (Table S5). If a CSS is reached (Figures 2c & 2d), inorganic nutrient is minimized through evolution but plant biomass and primary productivity are not systematically maximized nor minimized. Compare figure 2c and 2d. Evolution optimizes productivity when it comes at no costs in mineralization *α* (figure 2d), but such an optimization is not observed when such costs exist (figure 2c). When *α* is independent from *β* (figure 2d), the impact of evolution involves less dimensions (ie, impacts less compartments directly), so that this result confirms that evolution is more likely to be optimizing when the number of dimensions is reduced (Metz et al. 2008).

We stress that in CSS instances, final biomass and primary productivity always depends on basal biomass turnover *γ_0_*. (Table S4). In terms of management, it suggests that external disturbances (fire, pollution) not only directly impact ecosystem processes due to extra-mortality, but also further deteriorate their functional state by affecting evolutionary dynamics.

### Functional consequences of coupling aboveground and belowground traits

First, note that fitness (equation 12) is independent of belowground mineralization trait α. This happens because the nutrient consumption part of fitness only depends on the total amount of inorganic nutrient, as fixed by the resident population. Our model makes mean-field hypotheses, all mineral nutrient being equally accessible to all plant individuals. Differences in mineralization then add to the nutrient pool, at the advantage of any individual of the population, regardless of its phenotype. Changes in mineralization do not create any relative fitness difference. Consequently, existing links between *β* and *α* do not affect the selection of traits in our model. In spite of this conservative approach of mineralization effects, the coupling of aboveground and belowground processes still impacts the ecological consequences of evolution.

From an ecological point of view, the link between *β* and *α* modifies the plant biomass and productivity obtained through the evolutionary dynamics. Consider a model that would ignore the links between nutrient uptake *β* and mineralization *α*. From equation (10), it is easy to show that equilibrium plant biomass is then always increasing with *β* as *D^0^* increases with *β*. Linking *β* and *α* makes the variations more complex. If the relationship between *β* and *α* is positive, *D^0^* and *α* are both positively impacted by increases in *β* so that such evolutionary dynamics strongly increase expected plant biomass. Similarly, for situations in which the system settles at a given *β_CSS_*, if the relationship between uptake and mineralization is positive (*r>0*), then increasing this effect parameter *r* will in turn increase mineralization *α*, thereby increasing plant biomass and productivity (figure 3A). On the contrary, if the relationship is negative, the coupling between aboveground and belowground processes moderates the impacts of evolution on plant biomass and productivity or even reverses them (figure 4). Although the exact magnitude of change depends on parameter values, these results suggest that predictions that ignore links between growth rate and mineralization rate can be vastly misleading. Consequences may be far reaching: standing biomass and primary productivity largely affect ecosystem services and set the energetic basis and nutrient constraints for related food webs.

**Figure 3.**
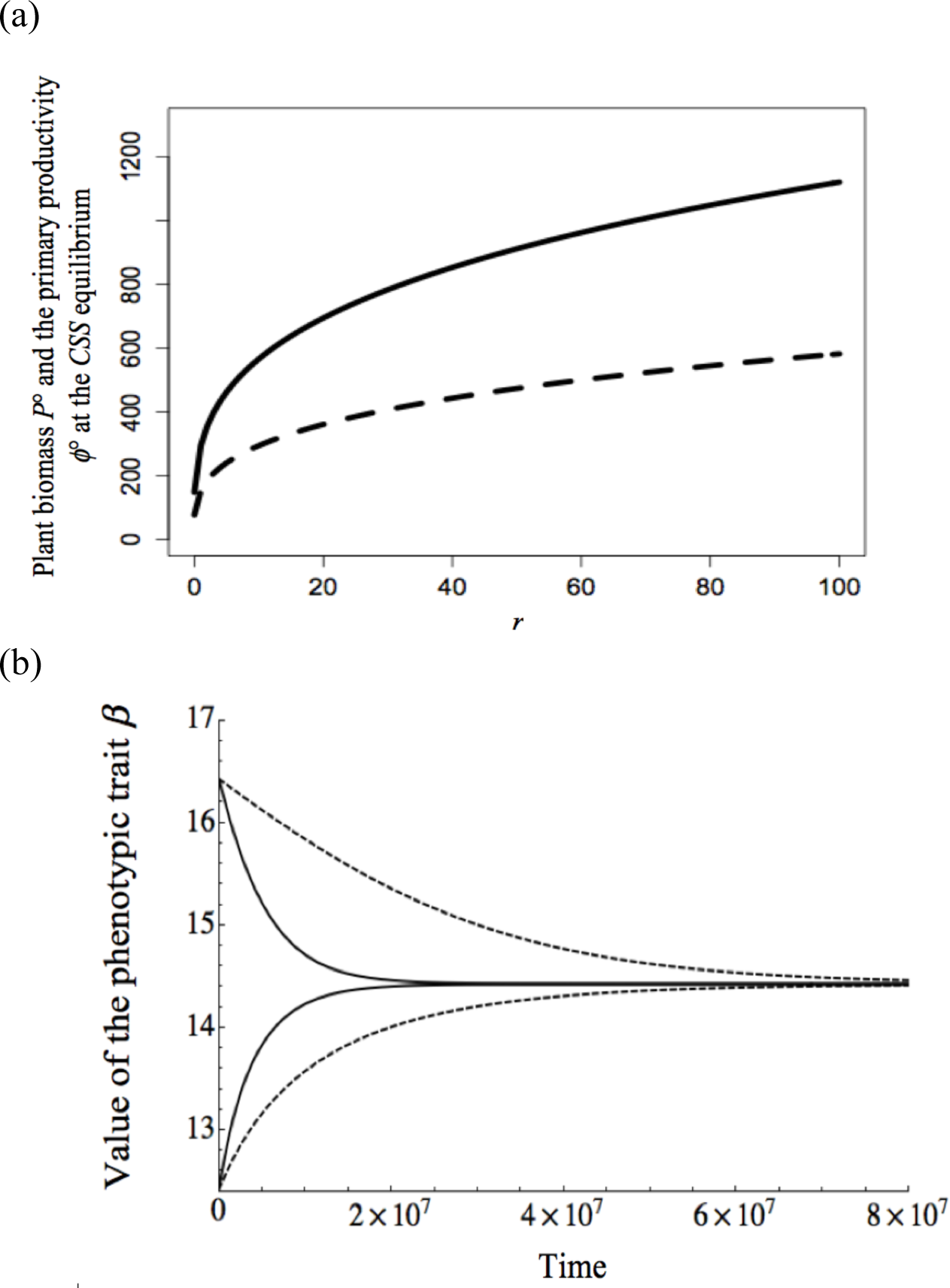
(a) Variation of the plant biomass *P*° (solid line) and the primary productivity *ϕ*° (dashed line) at the CSS depending on the strength of the impact of the evolution of *β* on belowground processes, *r*. *β* is fixed at the CSS value. A positive *α*-*β* relationship is assumed (eq 4b (Table 1)) (b) Variation of the evolutionary speed of *β* depending on whether coupling (dashed lines) exists or not (solid lines) between aboveground and belowground processes. *β_CSS_* is equal to 14.426. A negative *α-β* relationship is assumed (eq 4a ((Table 1))). Parameters values: *K*=1; *μ*=0.0001; *σ*=0.0001.

Many current works link the functioning of plant communities to their phenotypic states (Lavorel and Garnier 2002; Wright et al. 2004; Shipley et al. 2006). To understand the future functioning of ecosystems under disturbances, it may therefore be interesting to understand their stability in terms of phenotypic composition. To study this question, we analyze the “evolutionary resilience” of our system as measured by the return time to the initial phenotypic state following a disturbance. This measure of resilience is quite different from (but complementary to) the one classically used in ecology, as it is based on an analysis of evolutionary dynamics (trait variation) rather than on an analysis of the ecological equilibrium. On figure 3B, we show that this evolutionary resilience depends on the coupling between nutrient uptake and mineralization. This may be understood by accounting for changes in plant biomass observed in figure 3A.

Changes in plant biomass have important consequences for the pace of evolutionary dynamics, as larger plant populations lead to higher genetic variabilities. This is visible in equation (5), where the rate of change of the trait is linked to plant compartment size through the mutation process. Again, this has important, applied consequences. Consider a change in the phenotypic composition of plants. The return time to the evolutionary equilibrium (ie, evolutionary resilience) depends on the coupling between aboveground and belowground processes (Figure 3B). Consider an external disturbance that creates additional mortality, e.g., changing basal biomass turnover *γ_0_* therefore modifying *β_CSS_*. Depending on the strength of the link between aboveground and belowground processes, evolution toward the new evolutionary equilibrium may be fast or slow, hence affecting the robustness of ecosystem functioning. Here, the return time is much longer, as plant biomass is strongly reduced by a negative *β*-*α* relationship. We stress that the exact time associated with such evolutionary dynamics is generally unknown (it depends on the selective pressures, trade-off shape, genetic variability, generation time, etc), but the change in evolutionary resilience incurred by coupling aboveground and belowground processes is qualitatively robust.

In order to broaden the results illustrated by figure 3, we investigate how *r*, the impact of nutrient uptake *β* on mineralization *α*, affects plant biomass and evolutionary resilience (figure 4), compared to a reference scenario for which no impact exists (*r=0*). The left column assumes a negative impact (panels A & C), while the right column assumes a positive impact (panels B & D). As intuitively expected, when nutrient acquisition *β* and mineralization *α* are negatively correlated, plant biomass is decreased compared to the reference scenario (panel A). This is simply because evolutionary gain on one side (say, increase in nutrient uptake), is traded-off against nutrient availability on the other side (mineralization). Conversely, when the two traits are positively correlated, plant biomass is positively affected (panel B). We also show how such effects depend on two parameters of well-known functional importance: nutrient input (e.g., eutrophication), and basic turnover rate (e.g., fire, herbivory). Results show that the effects on plant biomass are exacerbated when nutrient input increases, or when basic turnover decreases. When the impact on mineralization allows for higher plant biomass (panel B & D), evolution is accelerated and the system more resilient (negative values on panel D: return time is reduced). Results illustrated by figure 4 clearly stress that to predict the effects of evolutionary dynamics on the functioning and resilience of the system one needs to know how aboveground and belowground processes are coupled. Such a link arguably depends on the species and ecosystem considered (table 1).

**Figure 4.**
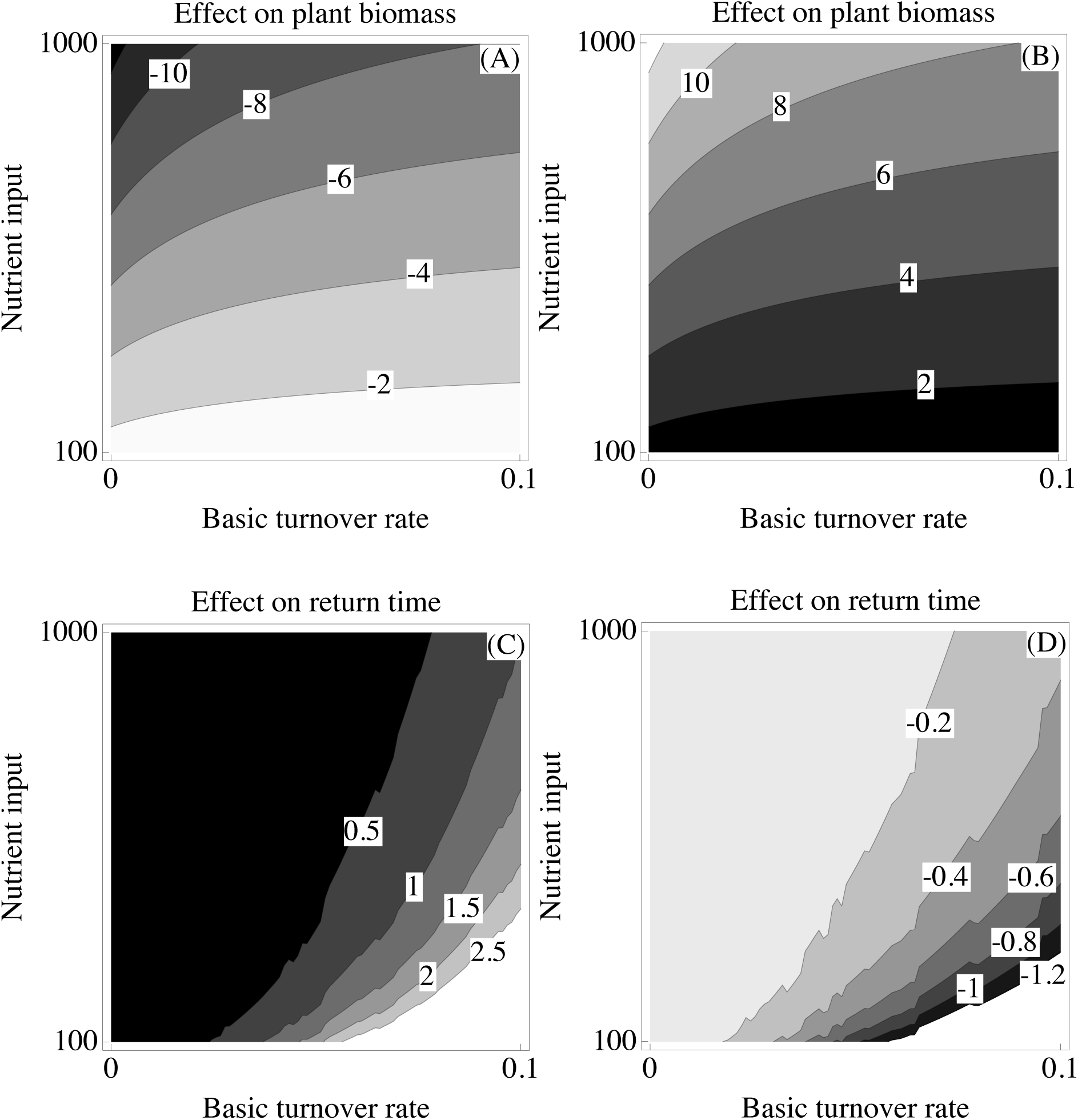
Variations in plant standing biomass (A, B) and resilience of the eco-evolutionary equilibrium (C,D) when nutrient input *I* and basic turnover rates *γ_0_* vary, depending on the impact of nutrient acquisition on mineralization rates (negative (see *eq 4a* (Table 1)): A & C; positive (see *eq 4b* (Table 1)): B & D). In each panel, variations are represented as differences with a reference no-coupling scenario where *r*=*0* (ie, effect on plant biomass ΔP=(P_r_)-(P_r=0_); effect on return time Δτ=(τ_r_)-(τ_r=0_)), P standing for the plant biomass at the evolutionary equilibrium, and τ for the return time to the evolutionary equilibrium following a disturbance of 5%. Lighter shades correspond to higher values. Numbers on contours in panels are expressed in thousand on panel A & C, and in million on panel B & D. Parameters values: *K*=1; *μ*=0.0001; *σ*=0.0001.

## Discussion

Consider a landscape made of multiple separated ecosystems, where local environmental conditions may involve changes in the trade-off shapes (Schluter 1995). Our results suggest that such variations in trade-off and variations in the coupling of belowground and aboveground processes strongly affect the functioning of the system. Ultimately, associated evolutionary dynamics can produce a range of behaviors, ranging from extinction to the intenance of a stable ecosystem, or even the unstable accumulation of plant biomass.

Regardless of the scenario, *N*° is minimized through evolution. This is in agreement the R* rule proposed by Tilman (1982). However, depending on the strength and the existence of trade-off constraints, three qualitatively different evolutionary outcomes have en identified: explosive R*, tragic R* and realized R* strategies. This last outcome corresponds to an evolutionary stable and convergent equilibrium where ecosystem nctioning critically depends on the coupling between aboveground and belowground processes. Also, we have proved that except for explosive R* strategies, plant biomass,imary productivity and dead organic matter are not necessarily maximized. In explosive R* strategies, the model cannot predict the future state of the system as another constraint will entually come into play (instead of the considered limiting nutrient). Such scenarios are nevertheless interesting, as they pinpoint cases in which evolution by itself may allow the stem to escape from one constraint to another, with important implications for predictions and management.

Even if the belowground trait *α* determining mineralization does not directly influence selected traits, it affects final biomass, productivity and evolutionary speed. When there is a strong link between aboveground and belowground processes, all aboveground phenotypic modifications cascade to constrain the energy allocated to belowground traits. Our results suggest that this can have important impacts on associated ecosystem services such as soil fertility or primary productivity. Several empirical works have suggested couplings between aboveground and belowground processes. For instance, evolution of plant defenses slows recycling processes (Grime et al. 1996; Whitham et al. 2003). Because the production of such defenses often incurs a cost in terms of growth (Herms and Mattson 1992), variations of defenses and their impacts on recycling relate well to the hypotheses of our model. A continental-scale study incorporating 13 vastly different ecosystems in North-America shows that investment in aboveground growth and biomass is linked to belowground processes such as the composition and mineralization activities of soil microbes (Zak et al. 1994). In spite of this increasing recognition that selections on aboveground and belowground traits are largely related, few evolutionary models incorporate this link explicitly (though see Reynolds and Pacala (1993); Gersani et al. (2001) for models of root competition and shoot/root ratios).

Our results have several important consequences in terms of conservation. First, local selection can decrease plant populations and negatively affect its role in the overall ecosystem functioning (in the case of tragic R* outcomes). Such outcomes occur when the benefit of an increase in nutrient uptake is constrained by a stronger cost in competition ability or survival. Such results are similar to those of other evolutionary models (Parvinen 2005; Boudsocq et al. 2011). Our study also gives new perspectives on existing links between plant evolution and ecosystem functioning. Runaway evolution occurs for concave trade-offs and population decreases (tragic scenarios) with convex trade-offs when inputs of nutrient are high and outputs are low. Evolutionary equilibrium is reached for convex trade-offs when inputs of nutrient are low and outputs are high. Even though an explicit test of these patterns is hardly possible because trade-off shapes are usually unknown, such dynamics correspond to contrasted situations that happen in nature and are usually considered separately (see also Boudsocq et al. (2011)).

It has often been postulated that evolution should maximize nutrient fluxes and increase primary productivity (e.g., Lotka 1922; Odum & Pinkerton 1955; Roff 1992). However, as shown by our study, taking into account unavoidable trade-offs and measuring fitness at the individual level, there is no reason to expect such effects. Some empirical studies that considered plant individual competition for resources have shown that primary productivity is not always maximized (e.g., Rankin,Bargum & Kokko 2007), in agreement with our results. Such negative relationships between community performance and individual competitiveness also have important implications for the improvement of crop yield potential in agricultural ecosystems (Denison 2012; Loeuille et al. 2013). Similarly, and contrary to predictions by Lotka (1922), aboveground and belowground selected strategies do not necessarily lead to tighter nutrient cycling.

Most works in plant community focus on either one trait or two traits linked by one trade-off function. An early example corresponds to the classical *r/K* theory, which organizes plant species along the growth/competitivity trade-off (Pianka 1970). Other examples include colonization/competition (Tilman 1994) or growth/defense trade-offs (Herms and Mattson 1992). Such a focus on one or two traits allows a degree of simplicity and a mechanistic understanding of eco-evolutionary dynamics (Boudsocq et al. 2011). A danger, however, is that it also tends to yield an adaptationist view of evolution that disregards the fact that individuals have many more traits, linked by multi-dimensional constraints (Gould and Lewontin 1979). Accounting for this complexity is a major challenge for evolutionary and community ecology. Here, we link four traits through allocation trade-offs. In spite of this added complexity and of the large number of trade-offs we tested, we have found some robustness in our results as only three qualitative functional outcomes have been identified. The multi-dimensional trade-off approach allows links with other multi-dimensional evolutionary theories of plant strategies (Grime 1977; Southwood 1988). For instance, some outcomes of our models (predicting a decrease in trait *β*), produce a syndrome of slow-growing conservative strategy very similar to stress-tolerant strategies introduced by Grime (1977).

In a context where databases of plant traits are systematically studied to predict the effects of global changes and the variations in ecosystem services (Lavorel and Garnier 2002), an important issue remains open. Plant species can be classified along a trade-off between acquisition and conservation of the resource (Díaz et al. 2004). Though this is clear for aboveground traits that define the leaf economic spectrum (Wright et al. 2004), such a spectrum for root traits is not fully clear. Mechanistic models could in the future help to predict (1) whether, (2) when evolution should lead to positive or negative correlations between aboveground and belowground traits and (3) the consequences for ecosystem functioning. Our result particularly suggests that resilience in trait composition strongly depends on how mineralization is linked to other traits such as nutrient uptake or turnover.

Because we focus on this issue of evolutionary multidimensionality, the ecological structure of our model is kept simple. Two extensions of this work would be particularly valuable. The first is related to the spatial context. In our model, the mean field hypothesis explains why mineralization is not present in the fitness definition we get from the present model. Accounting for spatial structures allows for a benefit of higher mineralization through local recycling (Barot et al. 2014) and nutrient compartment are then no longer minimized through evolution (Barot et al. 2015). Another extension is to account for other functional groups, as they crucially modify nutrient cycles. Herbivores affect nutrient spatial dynamics through dispersal at meta-ecosystem scale (McNaughton 1979). Gravel et al. (2010) have shown that spatial flows of material due to the nutrient diffusion or to plant or herbivore dispersal heavily impact the functioning of ecosystems. In fact, most instances of spatial flows of material involve higher trophic levels (e.g., McNaughton 1979; Helfield & Naiman 2001). In addition to changing nutrient constraints, incorporating higher trophic levels may constrain coexistence among plant phenotypes through apparent competition (P* rule, Holt et al. 1994).

Here we have shown how trade-offs between belowground (mineralization) and aboveground processes can determine ecosystem functioning in general, and more particularly plant productivity and ecosystem resilience. It would be relevant to test empirically the eco-evolutionary consequences of other trade-offs between belowground and aboveground functions. For example, by assessing the allocation of carbon and mineral resources to root exudates, roots, mycorrhizae and the aboveground system, or the allocation of belowground and aboveground defences against herbivores. Beyond the acknowledgment of various trade-offs, understanding such evolutionary dynamics, involving the coevolution of aboveground and belowground systems can profoundly change our view on the management of ecosystems.

## Acknowledgements

The authors are very grateful to Dominique Gravel and Nicolas Gross for their helpful and constructive comments on a previous version of the manuscript. T. Le Mao has been supported by a PhD grant from the French National Institute for Agricultural Research (INRA). The authors also acknowledge the support of CNRS and IRD.

